# The eIF2α kinase HRI triggers the autophagic clearance of cytosolic protein aggregates

**DOI:** 10.1101/2020.05.12.092072

**Authors:** Mena Abdel-Nour, Valeria Ramaglia, Athanasia A. Bianchi, Jessica Tsalikis, Hien N. Chau, Suneil K. Kalia, Lorraine V. Kalia, Jane-Jane Chen, Damien Arnoult, Jennifer L. Gommerman, Dana J. Philpott, Stephen E. Girardin

**Affiliations:** Department of Laboratory Medicine and Pathobiology, University of Toronto, Toronto, ON, Canada; Department of Immunology, University of Toronto, Toronto, ON, Canada; Krembil Research Institute, Toronto Western Hospital, University Health Network, Toronto, Canada; Institute of Medical Engineering & Science, MIT, Cambridge, MA, USA; INSERM U1197, Hôpital Paul Brousse, Bâtiment Lavoisier, 14 avenue Paul Vaillant Couturier, 94807 Villejuif Cedex, France; Université Paris-Saclay, Paris, France

**Keywords:** HRI, Integrated Stress Response, alpha-synuclein, Parkinson’s disease, protein aggregation, eIF2alpha, proteasome, autophagy, protein misfolding

## Abstract

Large cytosolic protein aggregates are removed by two main cellular processes, autophagy and the ubiquitin-proteasome system (UPS), and defective clearance of these protein aggregates results in proteotoxicity and cell death. Here we show that the eIF2α kinase HRI potentiates the autophagic clearance of cytosolic protein aggregates when the UPS is inhibited. In cells silenced for HRI, proteasome inhibition resulted in accumulation of aggresomes and ubiquitinated proteins, as well as cytotoxicity. Moreover, silencing of HRI resulted in cytotoxic accumulation of over-expressed α-synuclein, a protein known to aggregate in Parkinson’s disease, dementia with Lewy bodies, and multiple system atrophy. In agreement, protein aggregate accumulation and microglia activation were observed in the spinal cord white matter of 7-month old *Hri*^-/-^ mice as compared to *Hri*^+/+^ littermates. Moreover, aged *Hri*^-/-^ mice showed accumulation of misfolded α-synuclein, indicative of misfolded proteins, in the lateral collateral pathway, a region of the sacral spinal cord horn that receives visceral sensory afferents from the bladder and distal colon, a pathological feature common to α-synucleinopathies in humans where it may contribute to impaired micturition and/or constipation. Together, these results suggest that HRI contributes to a general cytosolic unfolded protein response (cUPR) that could be leveraged to bolster the clearance of cytotoxic protein aggregates.

---

Protein misfolding and aggregation are at the heart of neurodegenerative processes and can occur extracellularly as well as in the cytosol, the latter being evidenced by reported aggregation of various cytosolic proteins including α-synuclein (Parkinson’s disease and other synucleinopathies), tau (Alzheimer disease and tauopathies) and TDP-43 (Amyotrophic Lateral Sclerosis and Frontotemporal Dementia) to name a few ^1^. Central to the field of neurodegeneration is the idea that avoiding accumulation of these protein aggregates could prevent disease development and progression. Recent efforts have focused on understanding the mechanisms underlying the dynamic clearance of these aggregates, and in particular the role of autophagy and proteasome-mediated protein turnover in this process ^2,3^. However, how cells actually detect the initial formation of these aggregates to trigger their clearance is surprisingly poorly understood. Filling this gap would, most likely, greatly contribute to the design of novel therapeutic strategies in brain diseases. Indeed, understanding the underlying mechanisms accounting for the accumulation of, and the cell-to-cell propagation of, proteotoxic fibrillar aggregates, represent a major theme in multiple neurodegenerative diseases.

Heme-regulated kinase inhibitor (HRI), or eIF2α kinase 1, is one of the four eIF2α kinases, along with PERK, GCN2 and PKR, which collectively respond to a variety of cellular stresses and thereby trigger the Integrated Stress Response (ISR) ^4^. The main characteristics of the ISR are (i) the induction of a global translation shutoff directly caused by the phosphorylation of the translation initiation factor eIF2α and (ii) the upregulation of a stress-associated transcriptional reprograming dependent in part on the transcription factors ATF4 and ATF3. HRI was initially identified as a factor essential for controlling the expression of globin in red blood cells in response to changing levels of heme through a mechanism involving the dissociation of the chaperones Hsp9O and Hsc7O ^5,6^. However, HRI is ubiquitously expressed and induces eIF2α phosphorylation in various cell types in response to an array of stresses. In particular, the role of HRI in controlling the ISR in response to oxidative stress is well characterized ^7,8^. Indeed, phosphorylation of eIF2α and accumulation of eIF2α-dependent mRNA stress granules in cells treated with sodium arsenite, a potent oxidative stress inducer, is fully HRI-dependent. Whether HRI responds directly to the oxidative environment or to the consequences of the oxidative stress, such as accumulation of misfolded proteins, is currently unclear. However, the latter scenario is more likely given that HRI is also responsible for controlling eIF2α phosphorylation in response to heat shock ^9^ and proteasome inhibition ^10^, two conditions that lead to the accumulation of misfolded proteins in the cytosol.

In line with the proposed role of HRI in favoring cellular responses to proteotoxic stress and accumulation of misfolded proteins in the cytosol, we have recently demonstrated that the rapid engagement of several families of intracellular sensors of microbes, which are known to assemble into large molecular platforms (or signalosomes), require active control by an HRI-dependent pathway ^11^. This regulatory pathway, dependent on HRI, the heat shock protein HSPB8, eIF2α, ATF4 and ATF3, was coined the cytosolic unfolded protein response (cUPR), as it shares similarities with the pathway of regulation of protein folding in endoplasmic reticulum known as the unfolded protein response (UPR). In the absence of the cUPR, antimicrobial signalosomes are misfolded and display a defective capacity to trigger innate immune responses, such as pro-inflammatory signaling. This led to the more general suggestion that the cUPR in general, and HRI-dependent signaling in particular, could contribute to the homeostatic regulation of proteostasis in the cytosol, in part by inhibiting protein translation (following eIF2α phosphorylation) and by triggering a stress-dependent ATF3- and ATF4-dependent transcriptional reprogramming in response to the accumulation of cytosolic misfolded proteins.

In the context of neurodegenerative disorders, for which accumulation of cytotoxic protein aggregates is thought to play a major role, the potential contribution of HRI-dependent stress signaling remains uncharacterized. Interestingly, rare mutations in *EIF2AK1* and the related gene *EIF2AK2* (which encodes the eIF2α kinase PKR) are associated with developmental delay, white matter alterations, cognitive impairment and movement disorders ^12^. These observations are in line with previous reports that mutations in *EIF2B* are also associated with similar manifestations ^13^ and, more generally, that eIF2α is critical for neuronal health by integrating cellular stress pathways of the ISR ^14^. In agreement, it was recently observed that neuronal expression of HRI, although very low in resting conditions, was significantly increased following inhibition of protein degradation using a proteasome inhibitor, which resulted in constitutive inhibition of new protein synthesis through the HRI-eIF2α axis ^15^. This suggests that HRI plays a key role in neurons to maintain proteostasis, which could prevent accumulation of misfolded and potentially toxic proteins in these cells. Here, we provide evidence that HRI is critical for cytoprotection against proteotoxicity in cellular models of proteasome inhibition, likely by enhancing the autophagic degradation of protein cargos. Moreover, over-expression of α-synuclein, a protein known to be cleared by autophagy-mediated processes accumulated and was cytotoxic in HRI-silenced cells. In vivo, aged *Hri*^-/-^ mice displayed accumulation of protein aggregates and phospho-Sl29 α-synuclein in the central nervous system, supporting the notion that HRI-dependent cUPR may represent a novel mechanism that could be targeted to enhance clearance of protein aggregates in the context of neurodegenerative diseases.

## Experimental Procedures

### Cell lines and reagents

The human epithelial HeLa and HEK293T cell lines (American Type Culture Collection) were cultured in Dulbecco’s modified Eagle medium (DMEM) supplemented with 10% fetal calf serum (FCS), 2 mM L-glutamine, 50 IU penicillin, and 50 mg/ml streptomycin (Wisent Bio Products). Cells were maintained in 95% air and 5% CO2 at 37°C. Endotoxin-free FCS and phosphate-buffered saline (PBS) were from Wisent (Saint-Bruno-de-Montarville, Quebec, Canada).

Lentiviral knockdown of HRI expression was performed as previously described ^11^ and *HRI*^-/-^ cell line generation and characterization was also previously described ^11^. Stimulation with TNF (10 ng/ml, Cell Signaling technology) plus cycloheximide (10 μg/ml) was performed similar to previously ^16^. MG132 and chloroquine were from Sigma.

### Mice

*Hri*^+/-^ breeder mice were obtained from a previous study ^17^. Mice were bred and housed under SPF conditions at the Center for Cellular and Biomolecular Research, University of Toronto. All experiments using *Hri*^+/+^ and *Hri*^-/-^ mice were performed with 7 month old littermate mice generated from *Hri*^+/-^ x *Hri*^+/-^ cross. All mice experiments were approved by the Animal Ethics Review Committee of the University of Toronto. Genotyping was performed using primers described in a previous publication ^17^.

### Tissue collection

Mice whose spinal columns were harvested for histology were euthanized with CO_2_ and intracardially perfused with PBS, using a peristaltic pump. The spinal columns and brains were excised and post-fixed in 10% buffered formalin for 1 week prior to being processed into paraffin.

### Histology and immunohistochemistry

Seven micron paraffin coronal sections of mouse spinal cord and brain were mounted on Superfrost Plus glass slides (Knittel Glass, Germany) and dried in the oven (Precision compact oven, Thermo Scientific) overnight at 37°C. Paraffin sections were deparaffinated in xylene and rehydrated through a series of ethyl alcohol solutions.

Histology was performed using standard Hematoxylin & Eosin (H&E) and Luxol Fast Blue (LFB) stains to visualize inflammation and demyelination, respectively. The sections were subsequently dehydrated through a series of ethyl alcohol solutions and then placed in xylene before being coverslipped with Entellan mounting media (Merck Millipore, USA).

For the detection of protein aggregates, deparaffinated and rehydrated spinal cord sections were incubated in 1% Thioflavin S solution (Sigma-Aldrich, USA) for 10 minutes at RT. The sections were subsequently dehydrated through a series of ethyl alcohol solutions and then placed in xylene before being coverslipped with Entellan mounting media. Thioflavin S-bound protein aggregates were visualized under the microscope (Axio Imager Z1, Zeiss), using the EGFP filter (wavelength, 488 nm). Brain sections from donors with Alzheimer’s disease were used as positive controls.

For the immunohistochemistry, deparaffinated and rehydrated spinal cord sections were incubated in 0.3% H_2_O_2_ in methanol for 20 minutes to block endogenous peroxidase activity. Epitopes were exposed by heat-induced antigen retrieval in 10mM sodium citrate buffer (pH 6.0) in a pressure cooker placed inside a microwave set at high power (~800Watts) for 15 minutes. The non-specific binding of antibodies was blocked using 10% Normal Goat Serum (DAKO, Glostrup, Denmark) in PBS for 20 minutes at RT. Primary antibodies to detect either microglia/macrophages (rabbit monoclonal anti-Iba-1, clone EPR16589, abcam178847) or alpha synuclein phosphorylated on Ser129 (rabbit monoclonal anti-alpha-synuclein phospho S129, clone EP1536Y, abcam ab51253) were diluted (1:100) in Normal Antibody Diluent (Immunologic, Duiven, the Netherlands). Sections were incubated in the primary antibody solution overnight at 4°C. The following day, sections were incubated with the Post Antibody Blocking Solution for monoclonal antibodies (Immunologic) diluted 1:1 in PBS for 15 minutes at RT. Detection was performed by incubating the sections in the secondary Poly-HRP (horseradish peroxidase)-goat anti-mouse/rabbit/rat IgG (Immunologic) antibodies diluted 1:1 in PBS for 30 minutes at RT followed by incubation in DAB (3,3-diaminobenzidine tetrahydrochloride; Vector Laboratories, Burlingame, CA, U.S.A.) as chromogen. Counterstaining was performed with hematoxylin (Sigma-Aldrich) for 10 minutes. The sections were subsequently dehydrated through a series of ethyl alcohol solutions and then placed in xylene before being coverslipped with Entellan mounting media. Sections stained with secondary antibody alone were included as negative controls.

### Image analysis and quantification

For all staining, four to seven spinal cord sections from each of six *Hri*^+/+^ and six *Hri*^-/-^ mice were scored. The H&E and LFB stains of brain and spinal cord sections were screened for evidence of inflammation or demyelination, respectively, using a light microscope (Axioscope, Zeiss) connected to a digital camera (AxioCam MRc, Zeiss) and the Zen pro 2.0 imaging software (Zeiss). Since no signs of inflammation or demyelination were observed, no further quantification analysis was performed. The iba-1 immunostaining of brain and spinal cord sections was screened for evidence of microglial/macrophages clusters, as previously described ^18^, using a light microscope (Axioscope, Zeiss) connected to a digital camera (AxioCam MRc, Zeiss) and the Zen pro 2.0 imaging software (Zeiss). Since microglial/macrophages clusters were observed in the spinal cord but not in the brain, no further quantification analysis of iba-1 immunostaining in the brain was performed.

The Thioflavin S stain of brain and spinal cord sections was screened for evidence of Thioflavin S-bound protein aggregates, using the EGFP filter (wavelength, 488 nm) in a microscope (Axio Imager Z1, Zeiss) connected to a digital camera (AxioCam 506 mono, Zeiss) and the Zen pro 2.0 imaging software (Zeiss). Since Thioflavin S-bound protein aggregates were observed in the spinal cord but not in the brain, no further quantification analysis of iba-1 immunostaining in the brain was performed.

Quantitative analysis of the iba-1 immunostaining and the Thioflavin S staining of spinal cord sections was performed on 4x and 20x magnification, using ImageJ 1.15s (National Institute of Health, USA) imaging software. Images were calibrated in ImageJ. The white or grey matter areas of the spinal cord were measured at 4x magnification using the ImageJ freehand selections tool. The number of microglial/macrophages clusters and the number of Thioflavin S-bound protein aggregates was counted at 20x magnification over the entire white matter or grey matter areas of the spinal cord. Staining is expressed as number of microglial/macrophages clusters or the number of Thioflavin S-bound protein aggregates per mm^2^ of white matter or grey matter areas.

### Western blots

Unless otherwise indicated cells were lysed using RIPA buffer supplemented with protease and phosphatase inhibitors on ice. Lysates were then centrifuged at 13,000 x g to separate out membrane fractions / insoluble fractions and cytoplasmic proteins /soluble fractions. The membrane fractions were resuspended in RIPA buffer with Laemmli blue loading buffer and boiled to obtain the insoluble fractions.

Samples were then ran on acrylamide gels and transferred onto PVDF for blotting. For blotting of aggregates, membranes were first fixed for 30 minutes with 0.4% PFA. Membranes were then blocked with 5% milk or BSA in TBST and incubated with the indicated antibodies: mouse monoclonal anti-Tubulin (#T5168, Sigma, 1/10,000 dilution), rabbit anti-cleaved PARP (#9546, Cell Signaling, 1/1,000), rabbit anti-GFP (#ab290, Abcam, 1/5,000), rabbit anti-Hsc70 (#ab51052, Abcam, 1/1,000), mouse anti-α-synuclein (#ab1903, Abcam 1/3000), Ubiquitin, (Millipore, 1/200), rabbit anti-α-synuclein phospho-S129 (#ab51253, Abcam 1/3000).

### Co-Immunoprecipitation

For co-immunoprecipitation experiments cells were lysed using RIPA buffer supplemented with protease and phosphatase inhibitors on ice and subsequently centrifuged at 13,000 X G for 3 minutes. Protein G beads (Pierce) were washed twice with RIPA buffer and centrifuged at 2000 x G for 10 minutes before adding lysate and the indicated antibodies. The immunoprecipitation reaction was incubated overnight at 4°C. The following day samples were centrifuged at 10,000 X G for 5 minutes and washed 3 times with RIPA buffer supplemented with protease and phosphatase inhibitors before finally being resuspended in 120 μL of RIPA buffer with Laemmli blue loading buffer and boiled for 10 minutes before samples were subjected to SDS-PAGE.

### Immunofluorescence

HeLa cells were seeded onto glass coverslips (Warner Instruments) placed in 24-well plates at a density of 1.2 x10^5^. Following stimulation cells were fixed with 4% paraformaldehyde for 15 minutes. Coverslips were then washed twice with PBS and then permeabilized with 0.1% Triton in PBS for 10 minutes at room temperature. Samples were then washed two times with PBS and blocked for half an hour with blocking buffer (1% BSA in PBS).

Coverslips were then incubated with the indicated antibodies for 30 minutes in blocking buffer and washed twice with PBS in between antibody incubations. Antibodies were used as follows: Ubiquitin, 1:200 (Millipore), p62, 1:200 (Novus). Before mounting cells were stained with DAPI (4’,6-diamidino-2-phenylindole) dye (ThermoFisher) and mounted with Dako mounting medium and imaged by confocal microscopy and previously described ^19^.

### Transfections

For transient expression, transfections were performed with PEI MW25,000, and cells were lysed after overnight transfection. α-synuclein-GFP was obtained from Addgene. Plasmids encoding αsyn-L1 or αsyn-L2 have been described elsewhere ^20,21^.

### Luciferase assays

Luciferase assays were performed like previously described ^22^. HeLa cells were seeded in 24 well plates and transfected with 75 ng of Igκ luciferase reporter plasmid, 75ng of beta-Gal-SV40 expression plasmid and 300ng of pcDNA as a carrier per well, and the following day were used for infection experiments. NF-κB-dependent luciferase assays were performed in triplicate. Following infection cells were lysed in 100 μL Luciferase Lysis buffer (25mM Tris, 8mM MgCl_2_, 1% Triton, 15% glycerol). 10% of the lysate was added to darkened 96 well plates, and 100μL of Luciferase lysis buffer supplemented with 1mM DTT, 100μM ATP and Luciferin [1:500] was added to each sample and immediately read with a luminometer capable plate reader to measure luciferase activity. Beta-galactosidase activity was measured by adding 10μL of lysate to each well of a 96 well plate and 100μL of ONPG buffer (60mM Na_2_HPO_4_, 10mM KCl, 1mM MgSO_4_, 40mM NaH_2_PO_4_, 1mM DTT and ONPG 4mg/mL) was added to each well, incubated for half an hour and then read at 405 nM.

Luciferase assays to measure α-synuclein oligomerization were performd as previously described ^20,21^. HEK293T cells were transiently transfected in 6-well plates with Lipofectamine 2000 (Life Technologies), as per the manufacturer’s instructions. Plasmids transfected included αsyn-L1+αsyn-L2 or syn-L1+pcDNA combinations, with 1 μg of each plasmid used in each combination. At 24 hr post-transfection, the supernatant (1mL) was centrifuged at 3000xg for 5 min at 4°C. The cleared supernatant was used in the luciferase assay. The cells were scraped from each culture well in 1 mL PBS and then 50 μL of cells were transferred in triplicate to a 96-well, flatbottom, opague white plate (Greiner). The remaining cells were used for western blot analyses. Native coelenterazine (Prolume), a cell permeable substrate of Gaussia luciferase, was resuspended in nanofuel solvent (Prolume) to 5 mg/mL and dispensed per well to a final concentration of 20 mM by an automated plate reader (CLARIOstar, BMG Labtech). The bioluminescent signal generated by the luciferase enzyme was integrated over 2 sec before measurement at 480 nm. Cotransfection of a GFP vector was used to normalize for transfection efficiency in some experiments.

### Statistics

For all experiments statistic was performed using GraphPad Prism software 8.0 (GraphPad Software Inc, San Diego, CA, U.S.A.). Results of the experiments in cell lines were analyzed by Student t-test or ANOVA. For quantifications of thioflavin aggregates and microglial clusters, statistic was performed by multiple t test analysis and results were considered significant when P<0.05 at a 95% confidence level.

## Results

### HRI controls aggresome formation in response to proteasome inhibition

We silenced the expression of HRI in HeLa cells using small hairpin RNA directed against HRI sequence (shHRI) or a scramble sequence (SC). Cells were then treated for 4 hours with 5 μM MG132, a proteasome inhibitor and fixed cells were analyzed by immunofluorescence using antibodies against p62 and ubiquitin. As previously reported, proteasome inhibition using MG132 resulted in accumulation of p62+ and ubiquitin+ foci in a cytosolic area adjacent to the nucleus, known as the aggresome ^23^, in a small percentage of cells (Fig. 1A). Aggresomes result from the accumulation, at the microtubule organizing center (MTOC), of proteins cargos that are destined for degradation and migrated in a centripetal way along the microtubule network. Stimulation of the cells with higher doses of MG132 (up to 100 mM) and for longer periods of time (up to 16h) resulted in increased frequency and size of aggresomes, but this was associated with substantial cell death (data not shown). In HRI-silenced cells, we observed that both the frequency and architecture of MG132-induced aggresomes were dramatically affected. Aggresomes occurred more frequently in HRI knockdown (KD) cells (Fig. 1A-B) and also displayed larger diameter in average (1.84 μm +/-0.24 μm in HRI-silenced cells versus 1.35 μm +/-0.14 μm in control cells). Interestingly, while most small size aggresomes in control and HRI-silenced cells were both p62+ and ubiquitin+, large aggresomes (>2 μm in diameter), which were found nearly exclusively in HRI-silenced cells, were p62+/ubiquitin-(see examples of large aggregates in Field #1, Fig. 1A and quantifications in Fig. 1C). Moreover, proteasome inhibition, but not inhibition of autophagy using bafilomycin treatment, resulted in the accumulation of ubiquitinated proteins in HRI-silenced cells, as observed by western blotting on whole cellular extracts (Fig. 1D), suggesting that HRI silencing is in epistasis with autophagy rather than the UPS pathways.

**Figure 1:**
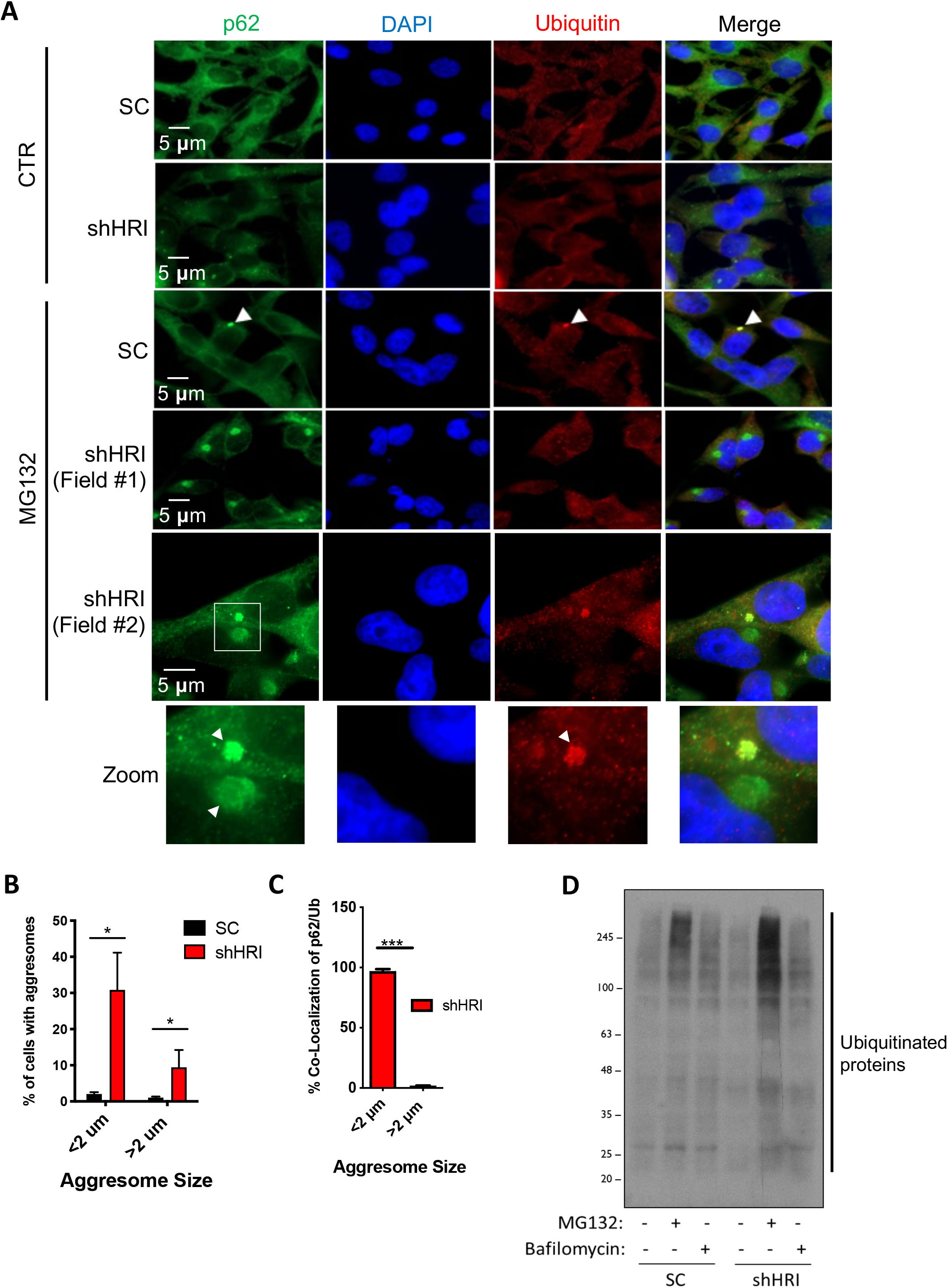
HRI controls aggresome formation in response to proteasome inhibition. **A-C**, HeLa cells transduced with lentiviral particles, targeting either a scrambled (SC) sequence or HRI (shHRI), were treated either with DMSO or with 5μM MG132 for 4 hours and stained with anti-p62 and anti-ubiquitin antibodies and analyzed by immunofluorescence (**A**) and quantified for the percentage of cells that have aggresomes larger and smaller than 2 μm (**B**) and for the frequency that these aggresomes have p62 and ubiquitin colocalization (**C**). **D**, HEK293T cells transduced with lentiviral particles targeting a scrambled sequence (SC) and HRI (shHRI) and treated for 4 hours with 10 μM MG132 and 10 nM Bafilomycin and analyzed by SDS PAGE using antibodies against ubiquitinated proteins. *, p<0.05; ***, p<0.001.

Together, these results suggest that, during cellular stress induced by the inhibition of the proteasome, HRI plays a critical role in controlling the dynamic targeting and flux of protein aggregates that are destined for degradation. In proteasome-inhibited cells, the absence of HRI resulted in the accumulation of aggresomes, the net build-up of ubiquitin-positive proteins in cells, and the formation of atypical and very large aggresomes that were p62+/ubiquitin-.

### HRI controls proteotoxicity in the face of proteasome inhibition through modulation of autophagy

Proteotoxicity and cell death are common features of the uncontrolled accumulation of protein aggregates. Treatment of HEK293T cells with 10 mM MG132 for 4 hours resulted in apoptotic cell death, as observed using western blotting by the accumulation of the cleaved form of PARP-1, and this cell death was exacerbated in shHRI cells (Fig. 2A). This effect was not caused by a general increase of sensitivity of shHRI cells to cell death because scramble-transduced and HRI-silenced cells were equally sensitive to cell death induced by tumor necrosis factor (TNF) plus cycloheximide (Fig. 2A). Similar sensitivity to MG132 was also observed in Crispr/Cas9-mediated HRI knockout cells as compared to wild type (WT) HEK293T cells (Fig. 2B). Importantly, shHRI cells appeared to be equally sensitive to autophagy inhibition by bafilomycin as control cells (Fig. 2C), thus showing that HRI silencing does not lead to a general sensitivity to the defective clearance of cytosolic cargos. Again, these results argue for the fact that HRI silencing is epistatic with autophagy pathways, which explains why HRI-silenced cells are particularly sensitive to the inhibition of the other arm of cytosolic protein clearance pathways, the UPS. In agreement, stimulation with 10 nM bafilomycin resulted in accumulation of the lipidated form of LC3 (LC3-II), a protein that is associated with autophagic cargos, and this accumulation was reduced in shHRI cells (Fig. 2D), suggesting that the homeostatic autophagic activity was consistently reduced in HRI-silenced cells. Similarly, the accumulation of p62 observed in bafilomycin-treated cells was reduced in shHRI cells (Fig. 2D). Therefore, these results together with those obtained in Figure 1, suggest a critical role for HRI in contributing to the autophagic arm of the cytosolic clearance of protein cargos.

**Figure 2:**
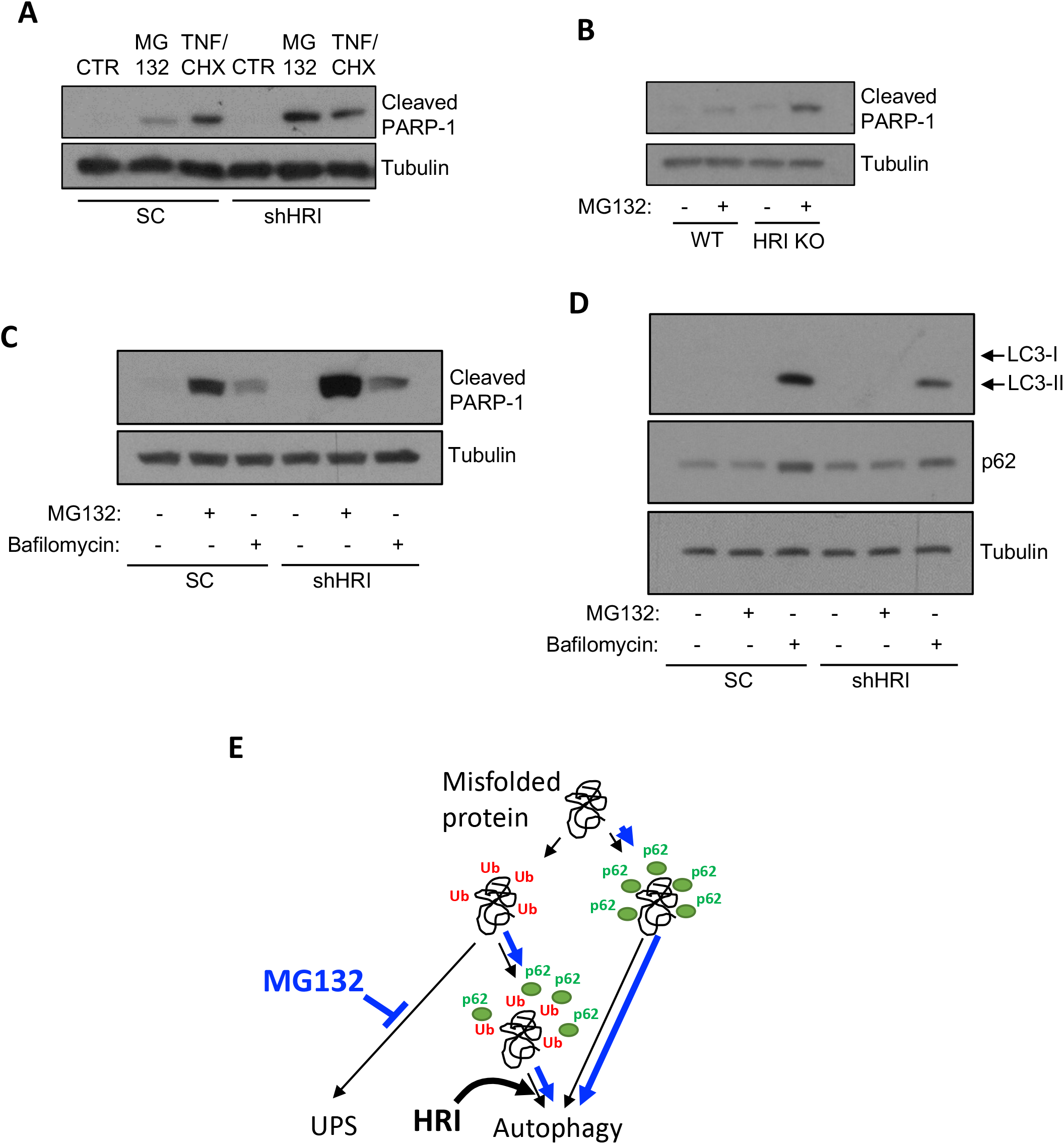
HRI controls proteotoxicity in the face of proteasome inhibition through modulation of autophagy. **A,** HEK293T cells transduced with lentiviral particles targeting a scrambled sequence (SC) and HRI (shHRI) were treated for 4 hours with 10 μM MG132 or TNF plus cycloheximide (CHX) and analyzed by SDS PAGE using antibodies against cleaved PARP-1 and tubulin. **B,** HEK293T HRI knockout (KO) or wild type (WT) cells were treated for 4 hours with 10 μM MG132 and analyzed by SDS PAGE using antibodies against cleaved PARP-1 and tubulin. **C-D,** HEK293T cells transduced with lentiviral particles targeting a scrambled sequence (SC) and HRI (shHRI) were treated for 4 hours with 10 μM MG132 and 10 nM Bafilomycin and analyzed by SDS PAGE using antibodies against cleaved PARP-1 and tubulin (C) or LC3, p62 and tubulin (D). **E**, schematic model representing the pathway(s) modulated by HRI that allow proteostasis.

### HRI controls responses to aggregating α-synuclein

α-synuclein, a protein that accumulates and forms aggregates in neurons in Parkinson’s disease and other synucleinopathies may impair the UPS early in the disease process ^24^ and is dominantly cleared from the cytosol by autophagy-mediated clearance ^25^. To monitor the role of HRI in controlling the accumulation of α-synuclein, we used a luciferase reporter system for which a split form of the luciferase enzyme is associated as two hemi-luciferase moieties in fusion with α-synuclein (αsyn L1 and αsyn L2). As α-synuclein self-assembles, the hemi-luciferases moiety are joined to generate a functional enzyme, and thus luciferase activity is a proxy of α-synuclein self-assembly ^20,21^. Over-expression of αsyn L1 and αsyn L2 resulted in significantly increased luciferase activity in lysates of shHRI cells as compared to scramble control cells (Fig. 3A, left), suggesting that a greater self-assembly of α-synuclein occurred in HRI-silenced cells. Interestingly, the cell culture medium prior to cell lysis also displayed greater luciferase activity in shHRI cells (Fig. 3A, right), and the difference was actually more pronounced than in the cell lysates. We reasoned that the accumulation of functional luciferase enzyme in the cell culture medium could be caused by cell death and the release of stable aggregates containing αsyn L1/αsyn L2. In agreement with this, we observed that over-expression of αsyn L1 and αsyn L2 was sufficient to cause apoptotic cell death, and this was exacerbated in HRI-silenced cells (Fig. 3B), suggesting that α-synuclein over-expression is particularly cytotoxic to shHRI cells. Finally, in HRI-silenced cells, over-expression of αsyn-GFP caused accumulation of multimeric forms of α-synuclein that were phosphorylated on Ser129 (Fig. 3C), which is a biochemical hallmark of α-synuclein aggregation ^26^. Moreover, these α-synuclein multimers were ubiquitinated, suggesting that the proteins are part of cargos that are destined for degradation, which build up in the cytosol of HRI-silenced cells (Fig. 3C). Interestingly, the accumulation of phospho-S129 and ubiquitin-positive GFP-αsyn was strongly enhanced after inhibition of the proteasome, but only in shHRI cells (Fig. 3C). We interpret these data to suggest that MG132-mediated inhibition of the proteasome imposes a stress on the autophagic machinery by shifting the normal homeostatic load of cargos needing to be cleared from the UPS to autophagy; as a consequence, the overload may not be fully absorbed by the autophagic machinery in HRI-silenced cells, in which autophagy may function below capacity (see Fig. 2), resulting in accumulation of phospho-S129 and ubiquitin-positive GFP-αsyn. Finally, it is worth noting that we consistently observed lower levels of over-expressed αsyn in the lysates of shHRI cells as compared to scramble control cells (see αsyn-GFP “input” western blot, Fig. 3C). We believe that this is caused by the fact that α-synuclein spontaneously aggregates in HRI-silenced cells, and thus disappears from the soluble fraction of our lysates. We consistently noted that greater amounts of α-synuclein could be retrieved in the RIPA-insoluble fraction (solubilized in urea) of shHRI cells, and that these forms of α-synuclein could only be blotted using an antibody against phospho-S129 synuclein, suggesting that the epitope recognized by the anti-α-synuclein was masked (data not shown). Together, these results suggest that HRI plays a key role in controlling the clearance and the cytotoxicity of α-synuclein, in cellular models of protein over-expression.

**Figure 3:**
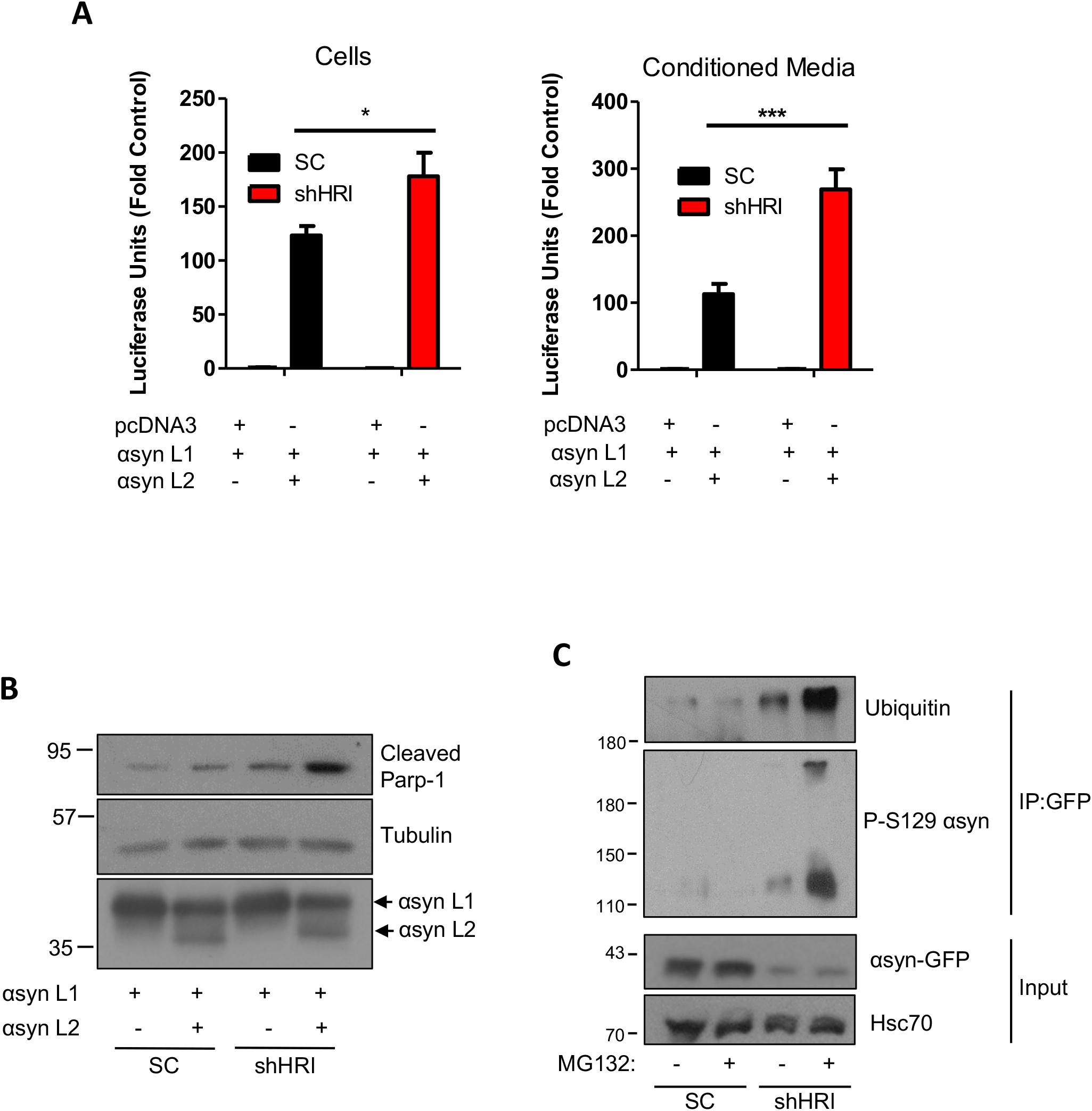
HRI controls responses to aggregating α-synuclein. **A**, HEK293T cells transduced with lentiviral particles targeting either a scrambled sequence (SC) or HRI (shHRI) were transfected with α-synuclein fused to a truncated luciferase (αsyn L1) and either pcDNA or α-synuclein fused to the remaining luciferase fragment (αsyn L2) and luciferase assays were performed on the cells (left) and the conditioned media (right). Bar graphs display a representative experiment with the means plotted from 2 experiments with 3 technical replicates each +/- S.D. (**, p<0.01). **B**, HEK293T cells transduced with lentiviral particles targeting either a scrambled sequence (SC) or HRI (shHRI) were transfected with αsyn L1 and either pcDNA or αsyn L2 and analyzed by Western blotting using the indicated antibodies. **C**, Coimmunoprecipitation assay using lysates of HEK293T cells transduced with lentiviral particles targeting a scrambled (SC) sequence or HRI (shHRI), transfected with GFP tagged α-synuclein following stimulation for 4 hours with 5μM MG132.

### Accumulation of protein aggregates in the central nervous system of aged *Hri*^-/-^ mice

In order to gain insights into the potential role of HRI in the control of protein aggregate clearance in a more physiological setting, we sought to determine if *Hri*^-/-^ mice would show signs of defective clearance in the central nervous system. To this end, we kept several pairs of littermate- and sex-matched *Hri*^-/-^ and *Hri*^+/+^ mice obtained from *Hri*^+/-^ intercrosses until the age of 7 months. At sacrifice, *Hri*^-/-^ and *Hri*^+/+^ littermate mice appeared overall healthy although we noted that all the knockout mice had developed splenomegaly (Fig. 4A), a characteristic that has been reported previously ^27,28^ in *Hri*^-/-^ mice, and relates to a progressive anemia and defective erythropoiesis in these mice. In contrast, brain weight was not significantly different between *Hri*^-/-^ and *Hri*^+/+^ littermate mice (Fig. 4B), and histology to detect myelin and inflammatory infiltrates in the brain (not shown) and spinal cord (Fig. 4C) did not show evidence of demyelination or overt inflammation in the central nervous system of *Hri*^-/-^ mice. Interestingly, clusters of Iba-1^+^ microglia/macrophages were observed in the spinal cord of *Hri*^-/-^ mice and were positive for Thioflavin S, a stain specific for amyloid-type protein aggregates (Fig. 4D). Quantification analysis revealed a significant increase in the density of Thioflavin S^+^ aggregates (Fig. 4E) and Iba-1^+^ clusters (Fig. 4F) in the spinal cord white matter of 7-month old *Hri*^-/-^ mice, suggesting that protein aggregates accumulate focally, resulting in local activation of the microglia. In support, we observed an upregulation of the protein CHOP, a marker of cellular stress typically associated with cell death, in the brains of 7-month old *Hri*^-/-^ mice (Fig. 4G). Together, these results suggest that deletion of Hri results in the spontaneous accumulation of protein aggregates, associated with cellular stress and microglial activation in the central nervous system of aged *Hri*^-/-^ mice.

**Figure 4:**
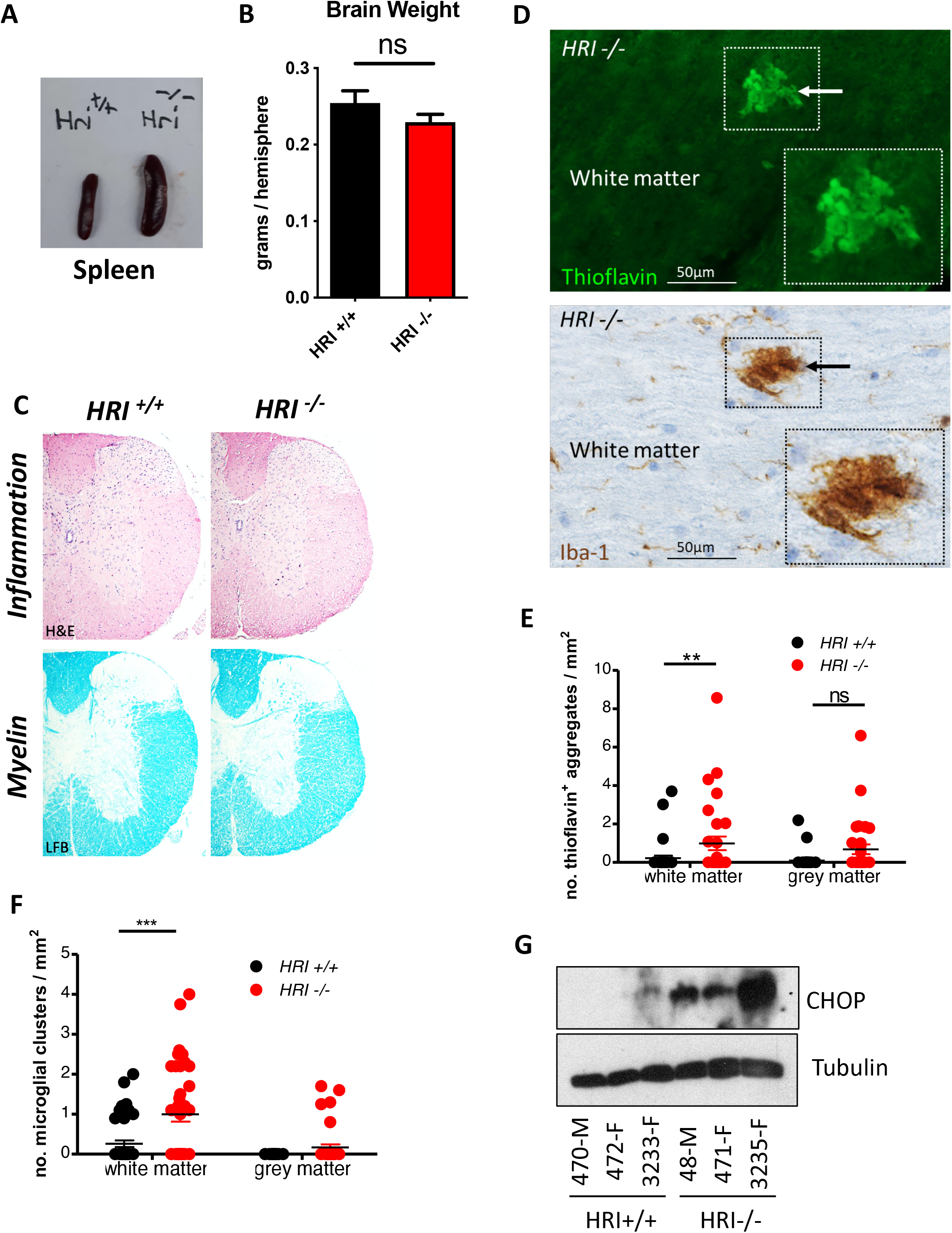
Accumulation of protein aggregates in the central nervous system of aged *Hri*^-/-^ mice. **A,** Splenomegaly in 7-month old *Hri*^-/-^ as compared to *Hri*^+/+^ littermates. **B**, Brain weight comparison between *Hri*^-/-^ as compared to *Hri*^+/+^ littermates (N=4 mice per group). **C**, Representative images of cross sections of spinal cord stained for hematoxylin & Eosin (H&E) to visualize inflammation and luxol fast blue (LFB) to visualize myelin in *Hri*^-/-^ as compared to *Hri*^+/+^ littermates. **D**, Representative images of longitudinal sections of spinal cord white matter stained for Thioflavin-S, to detect protein aggregates, or Iba-1, to detect microglia/macrophages. Images were from 2 adjacent sections, so that the same aggregate could be observed using both stains. Scale bar: 50μm. **E-F**, Quantification of Thioflavin-S^+^ (**E**) and Iba-1^+^ clusters (**F**) of the spinal cord white matter and grey matter from 7-month old wild-type (*Hri*^+/+^) and HRI knockout (*Hri*^-/-^) mice. G, Whole brain lysates from 7-month old *Hri*^-/-^ and *Hri*^+/+^ littermate mice (N=3 mice per group; M, male; F, female) analyzed by Western blotting using the indicated antibodies. For panels E and F, each dot represents the score from one spinal cord coronal section with 4-7 sections being scored per mouse. *, ** and ***, p < 0.05, 0.01 and 0.001, respectively.

### Accumulation of α-synuclein in the central nervous system of aged *Hri*^-/-^ mice

Given the results we have obtained using cellular systems, we next aimed to specifically determine if *Hri* deletion would affect α-synuclein in the mouse central nervous system. Western blot analysis of brain lysates from 7-month old *Hri*^-/-^ and *Hri*^+/+^ littermate mice revealed accumulation of S129 phospho α-synuclein dimers and oligomers (Fig. 5A) in *Hri*^-/-^ mice. While immunohistochemical analysis using an antibody against phospho S129 α-synuclein did not reveal specific areas of accumulation in the brains of *Hri*^-/-^ mice as compared to *Hri*^+/+^ mice (data not shown), 5/6 *Hri*^-/-^ but only 2/6 *Hri*^+/+^ mice displayed accumulation of phospho S129 α-synuclein in the lateral collateral pathway, a region of the sacral spinal cord horn that receives visceral sensory afferents from the bladder and distal colon (Fig. 5B). Together, these results suggest that *Hri* deficiency may lead to spontaneous accumulation of α-synuclein aggregates in the central nervous system in aged mice, specifically along the lateral collateral pathway that is known to be impaired in α-synucleinopathies in humans ^29^, supporting the notion that Hri-dependent pathways contribute to the clearance of potentially cytotoxic protein aggregates.

**Figure 5:**
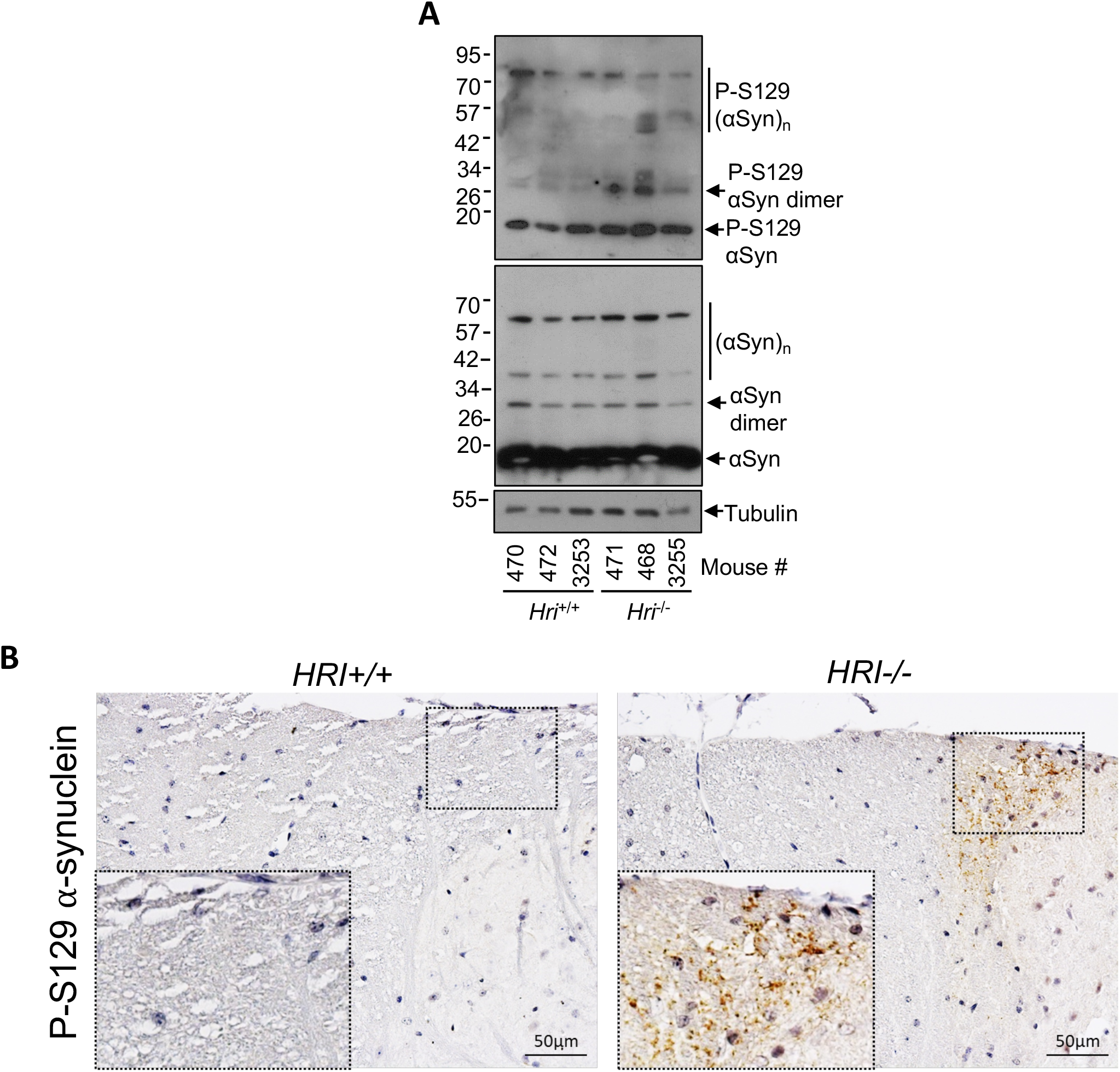
Accumulation of α-synuclein in the central nervous system of aged *Hri*^-/-^ mice. **A,** Whole brain lysates from 7-month old *Hri*^-/-^ and *Hri*^+/+^ littermate mice (N=3 mice per group) analyzed by Western blotting using the indicated antibodies. **B**, Representative images of wildtype and HRI knockout mice spinal cord coronal sections stained with anti P-S129 α-synuclein antibody. Scale bar: 50μm. Bar graphs represent the means +/- S.D. from 3-5 independent experiments.

## Discussion

The ISR is an evolutionary conserved stress response pathway that specifically controls the protein translation machinery via the targeting and phosphorylation of the translation initiation factor eIF2a. It is thus not surprising that the stresses detected by the ISR relate to protein homeostasis, such as amino acid starvation or endoplasmic reticulum stress caused by protein misfolding in this organelle. In this context, it is also not unexpected that HRI, one of the four mammalian kinases of the ISR, would participate in general protein homeostasis. Our previous work had reported a key role for HRI in the control of a cytosolic unfolded protein response or cUPR specifically following engagement of innate immune receptors that form large molecular platforms ^11^. Now, the present study shows that HRI is critical for cytoprotection against proteotoxicity in cellular models of proteasome inhibition, likely by enhancing the autophagic degradation of protein cargos, and over-expression of α-synuclein was cytotoxic in HRI-silenced cells. In agreement, we found that aged *Hri*^-/-^ mice displayed accumulation of protein aggregates and phospho-S129 α-synuclein in the central nervous system. We suggest that HRI plays a more general role in protein homeostasis by regulating the clearance of protein aggregates, likely through the upregulation of autophagy-dependent processes.

How HRI-dependent pathways contribute to an efficient turnover of protein aggregates through autophagy remains to be characterized. Based on our previous work, which demonstrated a key role for the heat shock protein (HSP) HSPB8 in HRI-dependent control of innate immune receptor scaffolds ^11^, it is conceivable that similar mechanisms involving HSPs contribute to the effects observed here. For instance, it is possible that the displacement of HSPs from HRI to cytosolic protein aggregates contributes to the activation of HRI, as previously suggested in the case of Hsc70 and Hsp90 ^27,30–33^. Then, ISR-dependent transcriptional reprogramming could result in the upregulation of specific HSPs that help in the autophagic clearance of protein cargos. In support for this, it is well characterized that HSPs play essential roles in assisting autophagy-dependent processes for the clearance of protein aggregates ^34^. A specific form of autophagy known as chaperone-assisted selective autophagy uses particular adaptors, such as the protein BAG-3, for the targeting of protein cargos to autophagosomes ^35^. Interestingly, this form of autophagy can specifically target p62-positive and ubiquitin-negative protein cargos, which could contribute to explain why p62-positive and ubiquitin-negative aggresomes accumulated in HRI-silenced cells in proteasome-inhibited cells (See Fig. 1). Also important to note is that BAG3-dependent chaperone-assisted selective autophagy typically uses the HSP, HSPB8 ^35^, which we previously found to be transcriptionally induced by HRI and involved in the cUPR ^11^. It is thus plausible that HSPB8, possibly together with a selection of other HSPs, would play a role in the general control of HRI-dependent proteostasis through autophagy. This may have implications for neurodegenerative disorders since autophagy, and more specifically BAG3- and HSPB8-dependent chaperone-assisted selective autophagy, has been shown to play important homeostatic functions in the brain ^36^.

The pathological analysis of the spinal cord from 7-month old *Hri*^-/-^ mice showed deposits of protein aggregates and clusters of microglia, reminiscent of pathological features found in the central nervous sytem of patients with neurodegenerative diseases ^37^. In addition, we found that in the 7-month old *Hri*^-/-^ mice α-synuclein accumulates in the lateral collateral pathway of the sacral spinal dorsal horn. This is a specialized region of the sacral spinal cord that receives visceral sensory afferents from the bladder and distal colon via the pelvic nerve ^38^. It is neurochemically distinct from other parts of the dorsal horn ^39,40^ and is well conserved among mammals ^38,41^. Notably, a pathological study on post-mortem spinal cord tissue has previously shown that α-synuclein is deposited at this region in individuals with α-synucleinopathies, including Parkinson’s disease, dementia with Lewy bodies, and multiple system atrophy, where it has been suggested to contribute to impaired micturition and/or constipation ^29^. Therefore deposition of α-synuclein in the lateral collateral pathway of the sacral spinal dorsal horn of aged *Hri*^-/-^ mice may be relevant to disease processes occurring in α-synucleinopathies in humans. Our data provide encouraging preliminary evidence that HRI-dependent control of the ISR may play a key homeostatic function in the central nervous system. It must be noted that these results were obtained without crossing our mice with transgenic animals that are typically used in neurodegenerative studies or using neurotoxic drugs, which can result in severe phenotypes but may be less physiologically relevant. Based on our results, it would be interesting to age *Hri*^-/-^ more than 7 months (up to 12-16 months) and also to perform behavioral studies to evaluate motor and cognitive impairment in these mice. With regards to the PD phenotype, a neuropathological analysis would be interesting to perform, focusing on sections of the spinal cord, brainstem (foremost the substantia nigra pars compacta of the midbrain vs the adjacent ventral tegmental area [an area of dopamine cells less vulnerable to PD]), hippocampus and cortex, as well as the peripheral nervous system which may be the initial site of onset of disease. While these assays are outside of the scope of the present initial study, we believe that our results provide an encouraging foundation for more in depth studies in *Hri*^-/-^ mice.

Importantly, our results suggest that the HRI-dependent pathway might represent a promising avenue for the treatment of neurodegenerative diseases or any pathology involving accumulation of cytotoxic protein aggregates. This approach might be more promising than targeting the more downstream event of eIF2α phosphorylation, which would have more dramatic side effects as it serves as a general hub for the multiple stresses of the ISR. Because the ISR and phosphorylation of eIF2α can be chronically upregulated in neurodegenerative diseases, attempts have been made to mitigate this pathway therapeutically ^42^. However, this may represent an overly simplistic approach that neglects the fact that if stress pathways are activated, it is because an underlying condition causes the stress in the first place. Moreover, targeting the messenger would not only be ineffective but could increase pathology, given that these pathways are induced to dampen the initial stress. Based on our results, we suggest that therapeutic interventions that would aim to transiently and locally amplify HRI-dependent signaling could be a viable option against the toxicity induced by cytosolic protein aggregates. A better understanding, at the biochemical level, of the cellular processes and pathways controlled by HRI will be beneficial for the rational design of such therapeutic strategies.

## Footnotes

This work was supported by grants from Canadian Institutes of Health Research (D.J.P and S.E.G). The authors declare no competing financial interests. We thank Drs Michael Schlossmacher and Julianna Tomlinson (University of Ottawa) for helpful discussions on α-synuclein biology along the way.

## REFERENCES

1. Hartl FU. Protein Misfolding Diseases. Annu Rev Biochem. 2017;86:21–26.

2. Ciechanover A, Kwon YT. Protein Quality Control by Molecular Chaperones in Neurodegeneration. Frontiers in neuroscience. 2017;11:185.

3. Sweeney P, Park H, Baumann M, et al. Protein misfolding in neurodegenerative diseases: implications and strategies. Translational neurodegeneration. 2017;6:6.

4. Pakos-Zebrucka K, Koryga I, Mnich K, Ljujic M, Samali A, Gorman AM. The integrated stress response. EMBO reports. 2016;17(10):1374–1395.

5. Chen JJ. Translational control by heme-regulated eIF2alpha kinase during erythropoiesis. Current opinion in hematology. 2014;21(3):172–178.

6. Chen JJ, Crosby JS, London IM. Regulation of heme-regulated eIF-2 alpha kinase and its expression in erythroid cells. Biochimie. 1994;76(8):761–769.

7. McEwen E, Kedersha N, Song B, et al. Heme-regulated inhibitor kinase-mediated phosphorylation of eukaryotic translation initiation factor 2 inhibits translation, induces stress granule formation, and mediates survival upon arsenite exposure. The Journal of biological chemistry. 2005;280(17):16925–16933.

8. Taniuchi S, Miyake M, Tsugawa K, Oyadomari M, Oyadomari S. Integrated stress response of vertebrates is regulated by four eIF2alpha kinases. Sci Rep. 2016;6:32886.

9. Lu L, Han AP, Chen JJ. Translation initiation control by heme-regulated eukaryotic initiation factor 2alpha kinase in erythroid cells under cytoplasmic stresses. Molecular and cellular biology. 2001;21(23):7971–7980.

10. Yerlikaya A, Kimball SR, Stanley BA. Phosphorylation of eIF2alpha in response to 26S proteasome inhibition is mediated by the haem-regulated inhibitor (HRI) kinase. The Biochemical journal. 2008;412(3):579–588.

11. Abdel-Nour M, Carneiro LAM, Downey J, et al. The heme-regulated inhibitor is a cytosolic sensor of protein misfolding that controls innate immune signaling. Science. 2019;365(6448).

12. Mao D, Reuter CM, Ruzhnikov MRZ, et al. De novo EIF2AK1 and EIF2AK2 Variants Are Associated with Developmental Delay, Leukoencephalopathy, and Neurologic Decompensation. Am J Hum Genet. 2020;106(4):570–583.

13. van der Knaap MS, van Berkel CG, Herms J, et al. eIF2B-related disorders: antenatal onset and involvement of multiple organs. Am J Hum Genet. 2003;73(5):1199–1207.

14. Bellato HM, Hajj GN. Translational control by eIF2α in neurons: Beyond the stress response. Cytoskeleton (Hoboken). 2016;73(10):551–565.

15. Alvarez-Castelao B, Tom Dieck S, Fusco CM, Donlin-Asp PG, Perez JD, Schuman EM. The switch-like expression of Heme-regulated kinase 1 mediates neuronal proteostasis following proteasome inhibition. Elife. 2020;9.

16. Killackey SA, Rahman MA, Soares F, et al. The mitochondrial Nod-like receptor NLRX1 modifies apoptosis through SARM1. Mol Cell Biochem. 2019;453(1-2):187–196.

17. Han AP, Yu C, Lu L, et al. Heme-regulated eIF2alpha kinase (HRI) is required for translational regulation and survival of erythroid precursors in iron deficiency. EMBOJ. 2001;20(23):6909–6918.

18. Michailidou I, Naessens DM, Hametner S, et al. Complement C3 on microglial clusters in multiple sclerosis occur in chronic but not acute disease: Implication for disease pathogenesis. Glia. 2017;65(2):264–277.

19. Tsalikis J, Tattoli I, Ling A, et al. Intracellular Bacterial Pathogens Trigger the Formation of U Small Nuclear RNA Bodies (U Bodies) through Metabolic Stress Induction. The Journal of biological chemistry. 2015;290(34):20904–20918.

20. Kalia LV, Kalia SK, Chau H, Lozano AM, Hyman BT, McLean PJ. Ubiquitinylation of α-synuclein by carboxyl terminus Hsp70-interacting protein (CHIP) is regulated by Bcl-2-associated athanogene 5 (BAG5). PLoS One. 2011;6(2):e14695.

21. Outeiro TF, Putcha P, Tetzlaff JE, et al. Formation of toxic oligomeric alpha-synuclein species in living cells. PLoS One. 2008;3(4):e1867.

22. Lee J, Tattoli I, Wojtal KA, Vavricka SR, Philpott DJ, Girardin SE. pH-dependent internalization of muramyl peptides from early endosomes enables Nod1 and Nod2 signaling. J Biol Chem. 2009;284(35):23818–23829.

23. Boyault C, Zhang Y, Fritah S, et al. HDAC6 controls major cell response pathways to cytotoxic accumulation of protein aggregates. Genes Dev. 2007;21(17):2172–2181.

24. McKinnon C, De Snoo ML, Gondard E, et al. Early-onset impairment of the ubiquitin-proteasome system in dopaminergic neurons caused by α-synuclein. Acta Neuropathol Commun. 2020;8(1):17.

25. Hou X, Watzlawik JO, Fiesel FC, Springer W. Autophagy in Parkinson’s Disease. J Mol Biol. 2020.

26. Hirai Y, Fujita SC, Iwatsubo T, Hasegawa M. Phosphorylated alpha-synuclein in normal mouse brain. FEBS Lett. 2004;572(1-3):227–232.

27. Liu S, Suragani RN, Wang F, et al. The function of heme-regulated eIF2alpha kinase in murine iron homeostasis and macrophage maturation. J Clin Invest. 2007;117(11):3296–3305.

28. Zhang S, Macias-Garcia A, Velazquez J, Paltrinieri E, Kaufman RJ, Chen JJ. HRI coordinates translation by eIF2αP and mTORC1 to mitigate ineffective erythropoiesis in mice during iron deficiency. Blood. 2018;131(4):450–461.

29. VanderHorst VG, Samardzic T, Saper CB, et al. α-Synuclein pathology accumulates in sacral spinal visceral sensory pathways. Ann Neurol. 2015;78(1):142–149.

30. Berwal SK, Bhatia V, Bendre A, Suresh CG, Chatterjee S, Pal JK. Activation of HRI is mediated by Hsp90 during stress through modulation of the HRI-Hsp90 complex. Int J Biol Macromol. 2018;118(Pt B):1604–1613.

31. Chen JJ, London IM. Regulation of protein synthesis by heme-regulated eIF-2 alpha kinase. Trends Biochem Sci. 1995;20(3):105–108.

32. Shao J, Grammatikakis N, Scroggins BT, et al. Hsp90 regulates p50(cdc37) function during the biogenesis of the activeconformation of the heme-regulated eIF2 alpha kinase. J Biol Chem. 2001;276(1):206–214.

33. Uma S, Hartson SD, Chen JJ, Matts RL. Hsp90 is obligatory for the heme-regulated eIF-2alpha kinase to acquire and maintain an activable conformation. J Biol Chem. 1997;272(17):11648–11656.

34. Dokladny K, Myers OB, Moseley PL. Heat shock response and autophagy-cooperation and control. Autophagy. 2015;11(2):200–213.

35. Klimek C, Kathage B, Wördehoff J, Höhfeld J. BAG3-mediated proteostasis at a glance. J Cell Sci. 2017;130(17):2781–2788.

36. Limanaqi F, Biagioni F, Gambardella S, Familiari P, Frati A, Fornai F. Promiscuous Roles of Autophagy and Proteasome in Neurodegenerative Proteinopathies. Int J Mol Sci. 2020;21(8).

37. Colonna M, Butovsky O. Microglia Function in the Central Nervous System During Health and Neurodegeneration. Annu Rev Immunol. 2017;35:441–468.

38. Morgan C, Nadelhaft I, de Groat WC. The distribution of visceral primary afferents from the pelvic nerve to Lissauer’s tract and the spinal gray matter and its relationship to the sacral parasympathetic nucleus. J Comp Neurol. 1981;201(3):415–440.

39. Anand P, Gibson SJ, McGregor GP, et al. A VIP-containing system concentrated in the lumbosacral region of human spinal cord. Nature. 1983;305(5930):143–145.

40. Kawatani M, Lowe IP, Nadelhaft I, Morgan C, De Groat WC. Vasoactive intestinal polypeptide in visceral afferent pathways to the sacral spinal cord of the cat. Neurosci Lett. 1983;42(3):311–316.

41. Nadelhaft I, Roppolo J, Morgan C, de Groat WC. Parasympathetic preganglionic neurons and visceral primary afferents in monkey sacral spinal cord revealed following application of horseradish peroxidase to pelvic nerve. J Comp Neurol. 1983;216(1):36–52.

42. Bond S, Lopez-Lloreda C, Gannon PJ, Akay-Espinoza C, Jordan-Sciutto KL. The Integrated Stress Response and Phosphorylated Eukaryotic Initiation Factor 2α in Neurodegeneration. J Neuropathol Exp Neurol. 2020;79(2):123–143.

